# The possible copepod link between kelp forests, the pelagic ecosystem and deep-sea carbon sequestration

**DOI:** 10.1101/2023.01.06.523004

**Authors:** Kristina Ø. Kvile, Marc Anglès d’Auriac, Dag Altin, Rolf Erik Olsen, Kasper Hancke

## Abstract

Kelp forests are dynamic coastal habitats that generate large amounts of carbon-rich detritus. The fate of this detritus is largely unknown and considered a missing link in global carbon budgets. Kelp detritus can serve as food for benthic invertebrates and pelagic invertebrate larvae, but we know close to nothing about the role of kelp detritus as food source for other zooplankton. Lipid-rich pelagic copepods constitute a key link from primary producers to higher trophic levels in marine boreal and arctic ecosystems, and they transport vast amounts of carbon into the deep sea. We conducted feeding experiments to test if the copepod *Calanus finmarchicus* can feed on fragments of two dominant kelp species, *Saccharina latissima* and *Laminaria hyperborea*. Such feeding would constitute an undescribed pathway from blue forests to pelagic consumers and deep-sea carbon sequestration. The experiment indicated that *C. finmarchicus* can ingest kelp particles, but the digestion is limited compared to a regular phytoplankton diet. Moreover, the results provide initial evidence that *L. hyperborea* contains substances that are toxic to copepods, an observation that warrants further research. Using specific qPCR assays to trace the consumption of kelp, we found that kelp DNA amplification signals were stronger for copepods fed with *S. latissima* than L. hyperborea, but we were not able to conclusively separate consumed kelp from DNA attached to the outside of the copepods.

## Introduction

Blue forests, coastal vegetated habitats such as kelp forests, seagrass beds and mangroves, are natural carbon sinks that contribute to regulating the global climate [1,2]. They store carbon in living biomass and deposit and sequester carbon in local and adjacent sediments or in the deep sea [3,4]. However, our knowledge of how this carbon is transported in the ocean is limited [5,6]. Comprehensive understanding of carbon fluxes is vital to quantify blue forests’ role in the carbon cycle and inform the discussion of whether the restoration and protection of blue forests may constitute a nature-based solution to mitigate climate change [1,2,7].

Kelps are among the most productive primary producers on the planet and form blue forests that dominate ∼1/4 of the world’s coastlines [8]. On average, >80% of the annual kelp biomass production is exported as detritus [9]. Detritus is generated through dislodgement of whole thalli or blades, or through erosion into fragments that range in size from small particles to larger pieces. Erosion occurs continuously and is likely more important than dislodgement during periods with normal wave conditions [9]. Moreover, dislodged blades will also erode, e.g., through grazing by sea urchins that turn kelp detritus into small fragments or fecal particles, increasing its availability to other consumers [10]. Kelp detritus may constitute a substantial part of the suspended particulate matter within and downstream of kelp forests [11,12] and is known to serve as a resource subsidy to other benthic habitats [9,13].

The kelp detritus is deposited at the sea floor or exported to adjacent or distant habitats, and small kelp particulates can be transported hundreds of kilometers [10]. The proportion that is exported is largely unknown and unaccounted for in global carbon budgets [4,7]. Recently, exported material of kelp and other macroalgae was documented across the global ocean, in surface and deep waters thousands of km from the coast [6], demonstrating that kelp organic matter is available in traceable amounts in coastal and ocean water masses, and might be an available food resource to zooplankton. Pelagic copepods of the genus *Calanus* play a key role in North Atlantic and Arctic pelagic ecosystems by converting carbon from primary producers into lipids available for higher trophic levels [14]. During the growth season, they store excess energy as lipids, and largely due to these energy-rich reserves, *Calanus* copepods are important prey for a range of animals. For example, *C. finmarchicus* dominates the zooplankton biomass throughout the North Atlantic Ocean and is the primary prey for large stocks of herring and mackerel and for larval cod [15]. *Calanus* copepods may descend for diapause at hundreds to >1000 m depth, a dormant state where they rely on the lipid reserves [16]. The copepods thereby transport carbon from the upper waters to the deep sea, a contribution to deep-sea carbon sequestration that is in the same order of magnitude as the passive sinking of organic material [17].

Fragmented kelp can serve as food for pelagic invertebrate larvae and various benthic invertebrates [18,19], but we have limited knowledge on the role of fragmented kelp as food for zooplankton. We hypothesized that pelagic copepods can consume fragmented kelp detritus and thereby form an undescribed link from blue forests to pelagic consumers and deep-sea carbon sequestration. To test this, we performed feeding experiments using the continuous *C. finmarchicus* culture at NTNU SeaLab (Trondheim, Norway). Beforehand, we collected material of the two dominant kelp species along the Norwegian coast, *Laminaria hyperborea* and *Saccharina latissima* [20], which was mechanically fragmented and stored in cool (4°C) and dark conditions to allow for bacterial degradation, aiming to mimic the natural decay process. During the feeding experiments, copepods were kept in bottles with filtered seawater mixed with either fresh or frozen kelp material, the phytoplankton used in the culture (*Rhodomonas baltica*) or no food. Supporting Table 1 gives an overview of the experimental setup.

## Results

### Change in particle number

As an indication of consumption, we compared the number of particles in the water before and after each experiment, focusing on particles in the 5-40 μm size range, where *C. finmarchicus* normally feeds [21,22]. We lack particle counts for Exp. 1-2, but the initial particle concentrations were consistent with Exp. 3-4. Exp. 3 and 4 lasted 24h, while experiment 5, with higher initial concentration, lasted 48h. Still, the water was replaced after 24h and the change in particle number measured after each 24h interval (Exp. 5A and 5B, respectively). Since the sample size per treatment was small (n≤6), the results should be interpreted with caution. We observed a decrease in particle numbers in all treatments (Figure 1), indicating feeding, except with no food, where there were few particles initially. The total volume of particles also generally decreased (Supporting Fig. 1). The ANOVA indicated no significant effect of treatment on mean change in particle number, nor of duration nor high/low initial particle concentration (Supporting Table 2). The number of particles <5 μm, which are smaller than what *C. finmarchicus* can efficiently filter [22], did not consistently decrease across treatments (Supporting Fig. 1).

**Figure 1:**
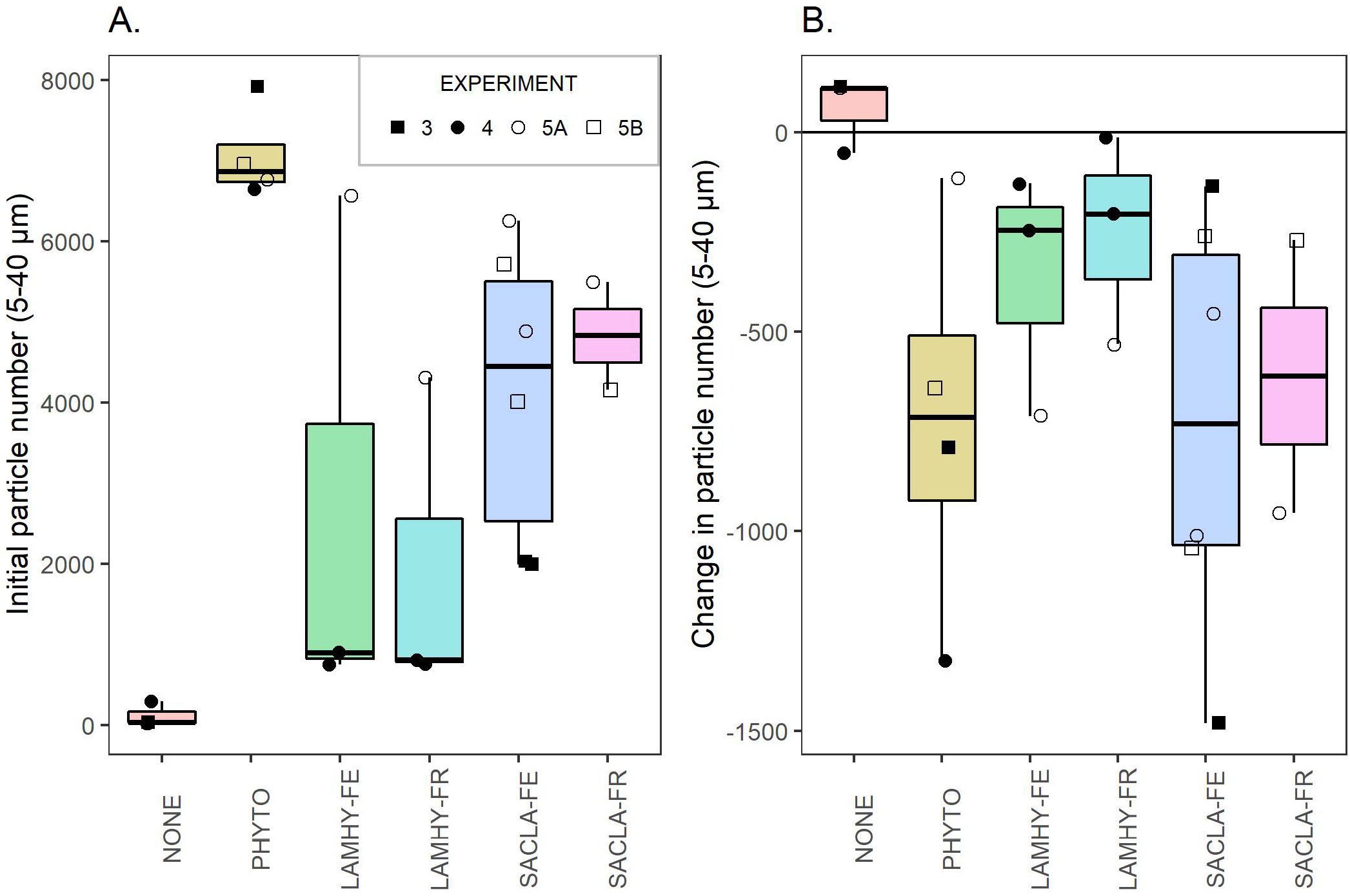
Initial number of particles in the water (A) and change in particle number during the experiments (B) for particles 5-40 µm, the normal feeding range of C. finmarchicus. Each point represents one observation (i.e., one experimental bottle measured before and after the experiment), with n per treatment ranging between 2 and 6, see Supporting Table 2 for details). The overlaying box plots show the median (line), the interquartile range (box), and 1.5 × the interquartile range (whiskers) per treatment. Unfilled symbols are observations from Exp. 5 with a higher initial volume of kelp than the other experiments. Food treatments: NONE: no food, PHYTO: phytoplankton (R. baltica); LAMHY-FE, LAMHY-FR: fresh or frozen L. hyperborea, respectively; SACLA-FE, SACLA-FR: fresh or frozen S. latissima, respectively.

### Fecal pellet production

Copepods fed with the phytoplankton diet they were used to in the culture produced a higher number of fecal pellets than copepods given kelp (Figure 2). For the phytoplankton treatment, the number of fecal pellets counted during a fixed 5 min interval ranged from 30 to 45 (i.e., 5-7.5 per copepod), but the total number was often higher. The ANOVA indicated significant differences between treatments, and the Tukey HSD post hoc tests confirmed that copepods fed phytoplankton produced more pellets than any other treatment, while the number produced by copepods fed kelp did not differ significantly from starved copepods (Supporting Table 3). When copepods had been exposed to the respective diets for 48 h (Exp. 5B), no additional pellets were observed for the *S. latissima* or ‘no food’ treatments (experiments with *L. hyperborea* terminated after 24h). This was reflected in a significant effect of duration in the ANOVA (Supporting Table 3).

### Mortality

In Exp. 2, we observed two dead and one sluggish copepod in one of two bottles with fresh *L. hyperborea* (no dead copepods were observed with frozen L. hyperborea). In Exp. 4, five out of six copepods in one of two bottles with frozen *L. hyperborea* were dead or sluggish (here, no dead copepods were observed in treatments with fresh L. hyperborea). In Exp. 5, when kelp concentrations were increased, all copepods exposed to fresh or frozen *L. hyperborea* were dead after 24h. Note that results from all experiments, including dead copepods, are displayed and can be distinguished in Figure 1 and Figure 2. We observed only one dead copepod among those fed frozen *S. latissima* (Exp. 1) or phytoplankton (Exp. 2), and no dead copepods in treatments with no food or fresh S. latissima.

**Figure 2:**
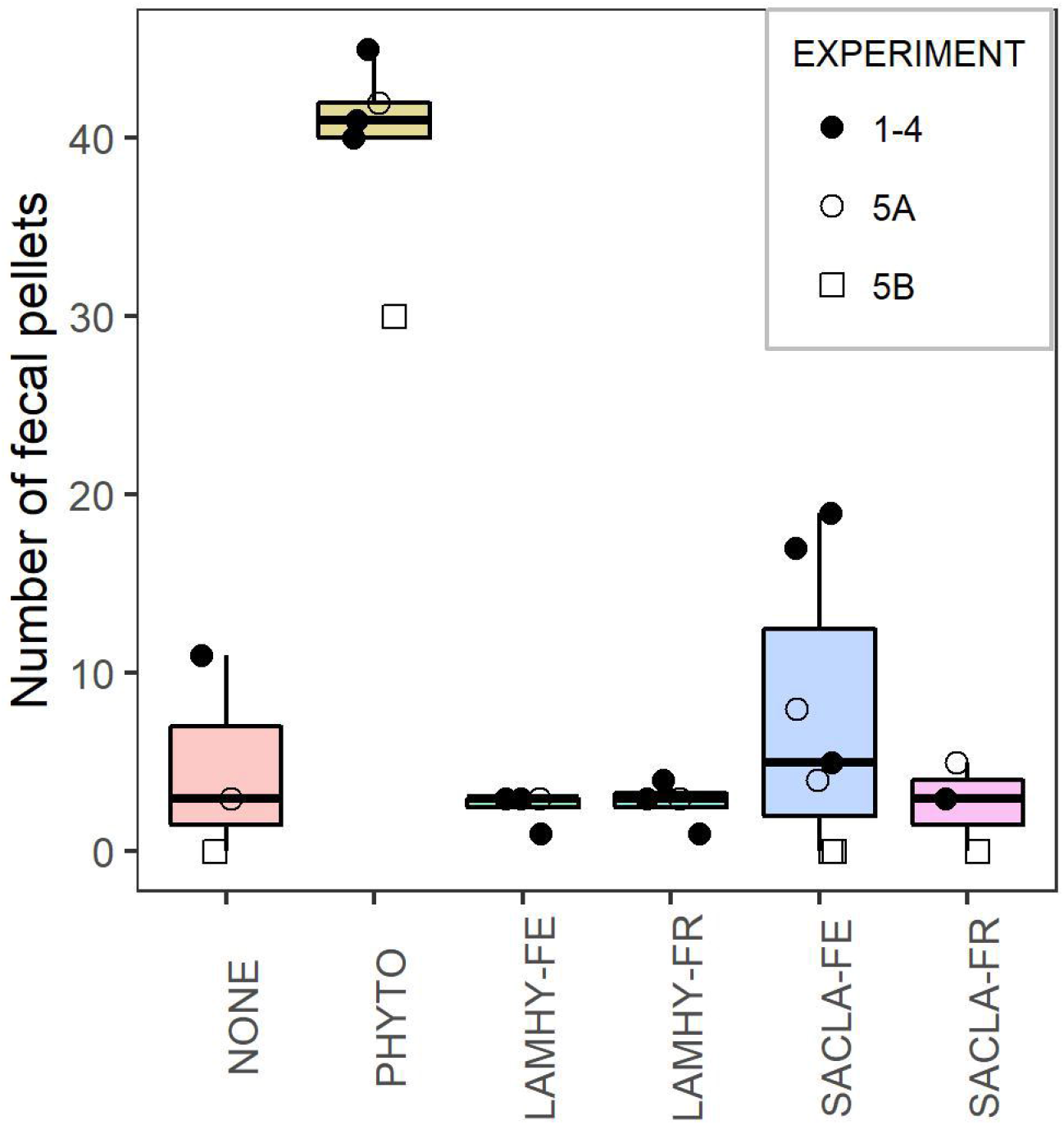
Number of fecal pellets counted after each experiment, using 5 min for searching per experimental bottle (for PHYTO, the total number of pellets was often higher). Each point represents one observation (i.e., one bottle), with n per treatment ranging between 3 and 7, see Supporting Table 3 for details. The overlaying box plots show the median (line), the interquartile range (box), and 1.5 × the interquartile range (whiskers) per treatment. Unfilled symbols are observations from Exp. 5 with a higher initial volume of kelp than the other experiments (Figure 1), and unfilled squares are from the second 24 h period of Exp. 5, after the copepods had been feeding on the respective diets for 24 h. Food treatments: NONE: no food, PHYTO: phytoplankton (R. baltica); LAMHY-FE, LAMHY-FR: fresh or frozen L. hyperborea, respectively; SACLA-FE, SACLA-FR, SACLA-P: fresh or frozen S. latissima, respectively.

### Presence of kelp DNA

We traced the presence of kelp DNA in copepods from the experiments using qPCR, first testing the quality of the different preservation and DNA extraction methods. Samples preserved in QuickExtract (QE) gave the strongest amplification of *C. finmarchicus* DNA (Supporting Fig. 2), and as using QE is the most time efficient method during extraction, we focused on these samples for the qPCR runs targeting kelp DNA (giving two copepod samples per experimental bottle). All qPCR assays performed optimally, with an efficiency of 100.4% and R^2^ of 0.992 for L. hyperborea, 103.2% and 0.971 for S. latissima, and 100.2% and 0.997 for C. finmarchicus. An optimal qPCR reaction doubles the target DNA product quantity at every amplification cycle, and the efficiency indicates how close the assay is to optimal amplification (100%). A qPCR run is considered satisfactory when the efficiency exceeds 90% [23].

As expected, when analyzing DNA extracted from copepods given no food or phytoplankton, there was no amplification of kelp DNA except for one replicate for the ‘no food’ treatment with the *L. hyperborea* assay (but the amplification was relatively week, Figure 3). The blanks similarly gave no kelp DNA amplification.

**Figure 3:**
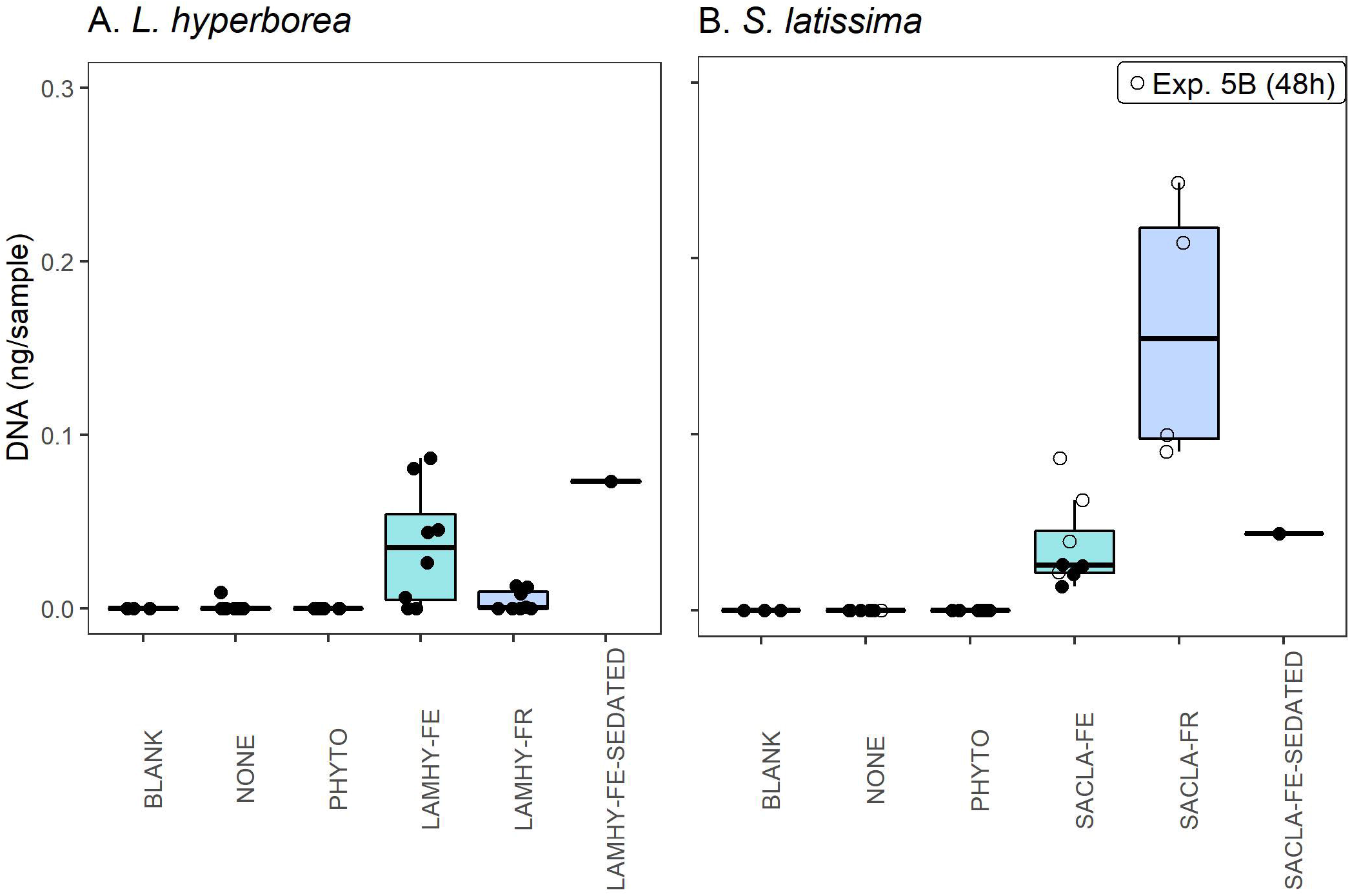
Estimated kelp DNA concentration (ng DNA/µl) extracted from copepods from different food treatments, compared to blanks (no sample material) and samples extracted from sedated copepods that were submerged in the kelp material for 1 min and then rinsed. Each point represents one observation (i.e., one copepod), with n per treatment ranging between 1 (sedated) and 8, see Supporting Tables 4 and 5 for details. The overlaying box plots show the median (line), the interquartile range (box), and 1.5 × the interquartile range (whiskers) per treatment. Unfilled circles in panel B are from Exp. 5B, after the copepods had been fed the respective diets for 48 h (treatments with L. hyperborea in Exp. 5 were terminated after 24h as the copepods were dead, and these samples were not analyzed). Treatments: BLANK: No sample; NONE: no food, PHYTO: phytoplankton (R. baltica); LAMHY-FE, LAMHY-FR: fresh or frozen L. hyperborea, respectively; SACLA-FE, SACLA-FR: fresh or frozen S. latissima, respectively; LAMHY-FE-SEDATED, SACLA-FE-SEDATED: copepods submerged in fresh L. hyperborea or S. latissima treatment while sedated, respectively.

All copepods fed fresh or frozen *S. latissima* showed target amplification for *S. latissima* DNA, with generally strongest amplification for samples fed with frozen material (Figure 3, Supporting Table 4). This could reflect higher consumption of frozen material, or the fact that these samples were from the 48h experiment. Kelp DNA concentrations also tended to be higher after 48h than 24h in copepods fed fresh *S. latissima* (Student’s t-test, t(−2.16) = 3.24, p = 0.11). Copepods fed *L. hyperborea* showed less consistent and generally weaker amplification than with S. latissima, with several samples not showing target amplification. The amplification was generally stronger for copepods fed with fresh than frozen L. hyperborea, the latter not differing significantly from treatments without kelp or blanks (Supporting Table 5).

However, copepods that had been sedated and submerged in kelp treatments for one minute (presumably not able to feed), and then rinsed with the same procedure as the other copepods, also showed kelp DNA amplification. For L. hyperborea, the amplification was of similar strength to the strongest amplification from samples from the experiments, whereas for S. latissima, several samples from the experiments showed stronger amplification, especially with the frozen material. Nevertheless, this indicates that the methodology could not separate kelp DNA attached to the outside of the copepods’ bodies from what was ingested.

## Discussion

We here suggest an undescribed pathway of carbon cycling from blue forests to pelagic consumers and deep-sea carbon sequestration, through the consumption of fragmented kelp detritus by pelagic copepods. We provide initial experimental results that we hope can trigger future studies on this potential trophic link, which is unaccounted for in carbon budgets and food web models and has implications for carbon sequestration and commercial fish stocks.

Kelp forests support a diverse ecosystem and numerous epifauna living on the kelp [24], and many fish species use this habitat for feeding and shelter [25]. However, the role of kelp forests for zooplankton is largely unexplored. *Calanus finmarchicus* is typically considered an oceanic species, with distribution across the northern North Atlantic Ocean [15]. But *Calanus* copepods are also found in coastal areas, and *C. finmarchicus* is transported in and out of Norwegian fjords seasonally [26]. Young saithe feeding above kelp forests along the Norwegian coastline predominantly fed on *C. finmarchicus* [25], illustrating the species’ potential as vector from kelp to higher trophic levels. Moreover, since macroalgae detritus is transported vast distances with ocean currents and present in the pelagic environment from the surface to >4000 m [6], we hypothesize that aged and fragmented kelp is an available food source for zooplankton in coastal and oceanic areas.

In our experiments, kelp detritus was available as particles in a suitable size range for *C. finmarchicus* [21,22]. The decline in particle concentrations in the 10-40 µm size range suggests kelp consumption by copepods (Figure 1), however, the low fecal pellet production indicates limited digestion (Figure 2). The decrease in particle number and the number of pellets produced tended to be higher for copepods fed *S. latissima* than L. hyperborea, but differences were not statistically significant and could be influenced by the higher mortality in *L. hyperborea* treatments. Moreover, we could not exclude that the copepods produced pellets based on feeding prior to the experiments, as the number of pellets from copepods fed with kelp and those given no food did not differ significantly, and no additional pellets were produced when copepods were fed *S. latissima* for more than 24 h.

The chemical composition of the kelp material might have impeded consumption. While kelps contain valuable nutrients such as omega-3 fatty acids and proteins, their composition differs from phytoplankton in several aspects. Kelps contain high fractions of carbohydrates, including components that are considered indigestible [27,28]. Kelps can also contain high levels of iodine, heavy metals and arsenic [27,28]. Aging and microbial degradation increases the kelp’s nutritional quality by reducing the C:N ratio and the concentrations of phlorotannins, which are known to deter grazing [9,19,29]. Experiments have shown that degradation for ∼1 week in seawater makes kelp fragments suitable as food for amphipods [19] and pelagic sea urchin larvae [18]. Also, deep-sea amphipods in situ preferred degraded kelp over fresh macroalgae [30]. Our results seem to suggest that *C. finmarchicus* is more likely to consume *S. latissima* than L. hyperborea. Possibly, *S. latissima* is more suitable as food, but its longer degradation time prior to the experiments (6 vs. 2 weeks) likely also influenced both its nutritional quality and concentrations of anti-grazing compounds. The high mortality in the treatments with *L. hyperborea* suggests that this material contained compounds that were toxic to the copepods. This could reflect its shorter degradation time, and contrasts with in situ studies concluding that detritus from *L. hyperborea* and other closely related species provides a valuable food source to benthic fauna communities [12,30–32].

Possibly, benthic detritivores and suspension feeders found in or adjacent to kelp forests may be better adapted to digest kelp detritus than pelagic zooplankton. Such adaptions may also be present in their pelagic stages, as sea urchin larvae can grow on kelp detritus [18]. Other zooplankton species might also be more likely to consume kelp detritus than C. finmarchicus. Although *Calanus* copepods are omnivores feeding on microzooplankton and phytoplankton [33], data on detritus feeding are conflicting [34]. In contrast, the krill *Thysanoessa inermis* in an Arctic fjord had an isotopic signature indicating >30% dietary contribution of kelp or rockweed carbon [possibly feeding close to the seafloor, 13], and the coastal copepod *Acartia tonsa* has been shown to ingest and incorporate brown algae (*Fucus*) material [while not being able to grow or survive on this alone, 35]. We may also speculate that the cultured copepods used here, which have lived >70 generations with continuous supply of a few phytoplankton species (*R. baltica, Dunaliella tertiolecta*), have lost their ability to digest or incorporate other food types. Cultured animals are likely to have less diverse gut microbiomes [36] and to differ genetically from their wild conspecifics [37].

We clearly detected kelp DNA in all copepod samples from the *S. latissima* treatments, while for L. hyperborea, the DNA amplification was weaker and less consistent, again suggesting that *C. finmarchicus* might more readily consume *S. latissima* than L. hyperborea. However, we also detected kelp DNA from copepods that had only been submerged in the kelp treatments without feeding, demonstrating that kelp material was attached to the outside of the copepods. The qPCR results can therefore not conclusively confirm that the copepods had consumed kelp. To our knowledge, the present study is the first to use species-specific qPCR assays to detect the consumption of macroalgae in pelagic copepods. DNA-based methods enable detecting specific species in animals’ diets [38], and do not suffer from the limitations of commonly used stable isotope methods, such as low taxonomic resolution and spatiotemporal variation in background isotope signatures. The qPCR protocols used here were recently developed and proved to efficiently trace kelp DNA in sediments and be robust to microbial degradation [39]. The use of qPCR assays can thus be an efficient method to detect consumption of kelp and other macroalgae by zooplankton and other animals. However, efficient cleaning procedures for body-surface DNA or delicate sampling techniques targeting stomach content are essential.

While detritus is unlikely to constitute a major part of pelagic copepods’ diet when more valuable food sources are available, such as during phytoplankton blooms, exported kelp could constitute a supplemental resource throughout the year and particularly when phytoplankton availability is low [13]. Based on our experiments, we cannot definitively conclude on the role of kelp detritus for zooplankton or if *Calanus* copepods can transfer kelp carbon to higher trophic levels or to the deep sea. We therefore recommend future studies with material from different macroalgae and zooplankton species, with the aim to elucidate the role of pelagic filter feeders as a pathway for carbon transport and turnover from blue forests.

## Materials and methods

### Preparation of kelp material

The kelp material was collected before the feeding experiments, when opportunities arose during field work to suitable areas. Therefore, the degradation period was longer (6 vs. 2 weeks) for *S. latissima* than L. hyperborea. *Saccharina latissima* was collected in the inner Oslofjord 2021-05-02 and kept cool (4°C) and dark in seawater that was renewed every second day during the first week. Thereafter, the material was blended in a kitchen blender (OBH Nordica Hero 6700, 1400W) in 4×30 seconds intervals, first mixing ∼0.3 g fresh kelp in 150 mL water. The blending was repeated five times (2021-05-11, 2021-05-14, 2021-05-20, 2021-06-03 and 2021-06-08, not adding new water or kelp). At the final day (2021-06-08), the material was wet-sieved at 60 µm and kept cool (i.e. fresh, SACLA-FE) or frozen (SACLA-FR) until the onset of the experiments (8-16 days later). We collected *L. hyperborea* in the outer Oslofjord 2021-06-01. The material was kept cool in ambient seawater, and blended with filtered seawater in the same manner as for *S. latissima* on two occasions (2021-06-03, 2021-06-08) before it was wet-sieved at 60 µm and kept cool (LAMHY-FE) or frozen (LAMHY-FR).

### Feeding experiments

Supporting Table 1 gives an overview of the experimental setup. In short, we conducted 5 experiments, i.e., different days with different combinations of treatments. Before each experiment, we randomly picked out *C. finmarchicus* individuals from the culture (almost exclusively adult females since this stage dominated in the culture), which were transferred to 1L bottles (6 ind./bottle) and placed in a plankton wheel with 0.8 rotations/min (keeping particles in suspension without disturbing the copepods). Each experiment included 4-7 bottles, giving 29 experimental bottles in total. The experiments typically included two replicates of either each SACLA treatment or each LAMHY treatment, one bottle with phytoplankton and one with no food, giving 3-6 replicates per treatment across experiments (Supporting Table 1). The experiments lasted 24h (Exp. 1-4) or 48h (Exp. 5). Food was added to the bottles beforehand, by mixing filtered (0.22 µm) seawater with either kelp material, the phytoplankton used in the culture (*Rhodomonas baltica*, PHYTO) or no food (NONE). In Exp. 5, the water was replaced and new food added after 24 h (Exp. 5B, Supporting Table 1). We aimed to feed the copepods ad libitum [>200 μg C L^-1^, 40]. We defined the volume of kelp or phytoplankton material to add to the bottles by equaling it to the volume of R. baltica normally given in the culture (7000 particles/mL, ∼1.33×10^6^ µm^3^/mL). Before each experiment, we estimated the number and volumes of particles in subsamples from the kelp and phytoplankton material using a cell counter (Multisizer3; Beckman Coulter Life Sciences, Indianapolis, IN, USA), and thereby calculated the volume of food to add (Supporting Fig. 1). In Exp. 5, we increased the amount of kelp by adjusting to the number, instead of volume, of phytoplankton normally given (which have a smaller mean size than the kelp particles). Before and after each experiment, we measured the particle density in 1 mL solution from a ∼20 mL water sample from the bottles using the cell counter.

After each experiment, all copepods were individually rinsed to remove any kelp material on the surface of the body by moving them with a pair of forceps through three baths of clean, filtered seawater. They were then placed in a drop of water, anesthetized by adding a drop of tricaine methanesulfonate solution (Finquel, 1.5 g/L seawater; Argent Laboratories, Redmond, WA, USA), and imaged using a CCD camera (Nikon DS-Fi1/U2, Tokyo, Japan) mounted on a Leica MZAPO stereomicroscope (Leica Microsystems, Wetzlar, Germany). They were immediately preserved individually in 0.5 mL ethanol (96%), 0.4 mL ATL buffer (QIAGEN, Germantown, MD, USA) or 0.2 mL QuickExtract (QE) DNA Extraction Solution (Lucigen, Middleton, WI, USA). The water in the experimental bottles was filtered at 35 µm to collect feces, which were picked out with a pipette and counted under the microscope, using a standardized time interval of 5 min per bottle.

To test if the washing procedure was sufficient to remove all kelp DNA on the outside surface of the copepods, we ran a separate experiment where copepods from the culture (that had not been in contact with kelp material) were completely anaesthetized and submerged for 1 min in fresh LAMHY (6 copepods) or SACLA (5 copepods) in the same dilution as used in Exp. 5. The copepods were then washed and preserved in the same way as in the experiments.

### DNA extraction and species-specific qPCR assay

We used a quantitative polymerase chain reaction (qPCR) assay with species-specific primers targeting *L. hyperborea* [39] and *S. latissima* (Anglès d’Auriac et al., in press) to detect kelp DNA in copepods. Samples were kept at 5°C from the experiments until Qpcr analyses (3-4 months), and we extracted DNA from whole single individuals. Samples preserved in ATL buffer or ethanol were extracted with the QIAGEN DNeasy Blood & Tissue kit. Samples preserved in QE were incubated for 6 min at 65.0°C followed by 2 min at 98.0°C.

We first tested the quality of DNA preservation and amplification from samples preserved in different solutions (ethanol, ATL buffer or QE), by running a *C. finmarchicus* species-specific qPCR protocol [41], i.e., targeting the copepod DNA. The protocol was modified after running a gradient qPCR identifying an optimal annealing temperature at 54°C (Supporting Fig. 2).

We ran separate qPCR plates for samples from *L. hyperborea* and *S. latissima* treatments, including samples from treatments with phytoplankton or no food in both plates (i.e., these were analyzed twice), as well as negative controls (blanks) and reference material of the respective kelp species. To ensure the presence of copepod material, all samples were analyzed for both *C. finmarchicus* and kelp DNA. Details of the amplification protocol are given in Supporting Table 6. All samples were replicated (two replicates for copepod samples, three replicates for blanks). Kelp reference material was collected in 2018 at 62.71°N/6.34°E and 62.78°N/6.48°E for *L. hyperborea* and S. latissima, respectively, and stored as extracted DNA in eluate at -20°C. Standard curves were constructed using serial dilutions of the reference material (ng/µL DNA) in triplicates. The DNA content per sample (ng DNA) was estimated from the standard curve, averaging over all replicates per sample after validating positive signals against the melt curve analysis. Standard curves were also used for calculating the efficiency of the qPCR assay. All qPCR assays were carried out using a Bio-Rad CFX96 instrument.

## Data analyses

We compared the change in particle number; the number of fecal pellets produced; and amplification of kelp DNA between treatments, using ANOVA followed by Tukey HSD *post hoc* tests. For the change in particle number and number of fecal pellets, the ANOVA also included effects of duration (48h in Exp. 5B, 24h in the others) and initial particle concentration (“low” in Exp. 1-4, “high” in Exp. 5). Since the sample size per treatment was small (n≤8), the results of the statistical tests should be interpreted with caution. All analyses and plots were made in R [42].

## Acknowledgements

We thank Rune Edvin Haldorsen and Lise Tveiten for help during the collection of kelp material, Iurgi Salaberria and Sarah B. Ørberg for support during the experiment and DNA analyses, and Dorte Krause-Jensen for valuable discussions and guidance.

## Supporting Information

Supporting tables and figures are compiled in the document SuppInfo_CopeLink_Kvile_2023.pdf.

