## Supporting Information for "The possible copepod link between kelp forests, the pelagic ecosystem and deep-sea carbon sequestration"

Supporting Table 1: Overview of the feeding experiments, with termination date, experiment number, experimental bottle ID and treatment. All experiments lasted for 24h, except Exp. 5, which lasted 48 h. For Exp. 5, the water was replaced and new food added after the first 24h and continued for another 24 h, but the treatments with *L. hyperborea* (5A) terminated after the first period. Food treatments: LAMHY-FE, LAMHY-FR: fresh or frozen *L. hyperborea*, respectively; SACLA-FE, SACLA-FR: fresh or frozen *S. latissima*, respectively; NONE: no food, PHYTO: phytoplankton (*R. baltica*).

| DATE | EXPERIMENT | BOTTLE | TREATMENT |
|---|---|---|---|
| 2021-06-17 | 1 | A | SACLA-FE |
| 2021-06-17 | 1 | B | SACLA-FR |
| 2021-06-17 | 1 | C | SACLA-FE |
| 2021-06-17 | 1 | D | SACLA-FR |
| 2021-06-17 | 1 | E | NONE |
| 2021-06-17 | 1 | F | PHYTO |
| 2021-06-18 | 2 | A | LAMHY-FR |
| 2021-06-18 | 2 | B | NONE |
| 2021-06-18 | 2 | C | LAMHY-FE |
| 2021-06-18 | 2 | D | LAMHY-FR |
| 2021-06-18 | 2 | E | PHYTO |
| 2021-06-18 | 2 | F | LAMHY-FE |
| 2021-06-22 | 3 | B | PHYTO |
| 2021-06-22 | 3 | D | SACLA-FE |
| 2021-06-22 | 3 | E | SACLA-FE |
| 2021-06-22 | 3 | F | NONE |
| 2021-06-23 | 4 | A | LAMHY-FE |
| 2021-06-23 | 4 | B | NONE |
| 2021-06-23 | 4 | C | LAMHY-FE |
| 2021-06-23 | 4 | D | PHYTO |
| 2021-06-23 | 4 | E | LAMHY-FR |
| 2021-06-23 | 4 | F | LAMHY-FR |
| 2021-06-24 | 5A | G | LAMHY-FE |
| 2021-06-24 | 5A | H | LAMHY-FR |
| 2021-06-25 | 5B | A | NONE |
| 2021-06-25 | 5B | B | SACLA-FE |
| 2021-06-25 | 5B | C | SACLA-FR |
| 2021-06-25 | 5B | D | SACLA-FE |
| 2021-06-25 | 5B | F | PHYTO |

Supporting Table 2: Effects of treatment, duration and intial food concentration on the mean change in particle concentration (5-40 µm) during the experiments. The upper part of the table shows results from a three-way ANOVA of the effects of treatment, experimental duration (48h in Exp. 5B, 24h in the others) and intial food particle concentration (*high* in Exp. 5 and *low* in the others) on change in particle number. 'SSn': Sum of Squares in the numerator (i.e. SS effect); 'SSd': Sum of Squares in the denominator (i.e. SS error); 'DFn': Degrees of Freedom effect; 'DFd': Degrees of Freedom error; 'F': F-value; 'p': p-value; 'p<.05': significance at the 0.05 level; 'ges': Generalized Eta-Squared measure of effect size.

The lower part of the table shows results from Tukey HSD *post hoc* assessing the significance of differences between pairs of treatments (the other two variables had only two levels). 'n1'/'n2': the number of observations in group1 and group2; 'estimate': difference in observed means in group1 and group2; 'conf.low'/'conf.high': lower and upper end points of the confidence interval around the difference; 'p.adj': p-value after adjustment for multiple comparisons; 'p.adj.signif': significance at the 0.05 level (ns: not significant). Food treatments: LAMHY-FE, LAMHY-FR: fresh or frozen *L. hyperborea*, respectively; SACLA-FE, SACLA-FR: fresh or frozen *S. latissima*, respectively; NONE: no food, PHYTO: phytoplankton (*R. baltica*).

| Effect | SSn | SSd | DFn | DFd | F | p | p<.05 | ges |
|---|---|---|---|---|---|---|---|---|
| TREATMENT | 1.8e+06 | 2,586,664 | 5 | 13 | 1.8e+00 | 0.19 | | 4.1e-01 |
| DURATION | 1.1e+05 | 2,586,664 | 1 | 13 | 5.7e-01 | 0.46 | | 4.2e-02 |
| INITIAL_CONCENTRATION | 1.8e+00 | 2,586,664 | 1 | 13 | 8.9e-06 | 1.00 | | 6.9e-07 |

| group1 | group2 | n1 | n2 | estimate | conf.low | conf.high | p.adj | p.adj.signif |
|---|---|---|---|---|---|---|---|---|
| LAMHY-FE | LAMHY-FR | 3 | 3 | 113 | -1,016 | 1,241 | 1.00 | ns |
| LAMHY-FE | SACLA-FE | 3 | 6 | -369 | -1,346 | 609 | 0.82 | ns |
| LAMHY-FE | SACLA-FR | 3 | 2 | -250 | -1,512 | 1,012 | 0.99 | ns |
| LAMHY-FR | SACLA-FE | 3 | 6 | -481 | -1,459 | 496 | 0.61 | ns |
| LAMHY-FR | SACLA-FR | 3 | 2 | -362 | -1,624 | 899 | 0.93 | ns |
| NONE | PHYTO | 3 | 4 | -776 | -1,831 | 280 | 0.22 | ns |
| NONE | LAMHY-FE | 3 | 3 | -420 | -1,549 | 709 | 0.83 | ns |
| NONE | LAMHY-FR | 3 | 3 | -307 | -1,436 | 821 | 0.94 | ns |
| NONE | SACLA-FE | 3 | 6 | -789 | -1,766 | 189 | 0.15 | ns |
| NONE | SACLA-FR | 3 | 2 | -670 | -1,932 | 592 | 0.54 | ns |
| PHYTO | LAMHY-FE | 4 | 3 | 356 | -700 | 1,411 | 0.88 | ns |
| PHYTO | LAMHY-FR | 4 | 3 | 468 | -588 | 1,524 | 0.70 | ns |
| PHYTO | SACLA-FE | 4 | 6 | -13 | -905 | 879 | 1.00 | ns |
| PHYTO | SACLA-FR | 4 | 2 | 106 | -1,091 | 1,303 | 1.00 | ns |
| SACLA-FE | SACLA-FR | 6 | 2 | 119 | -1,010 | 1,248 | 1.00 | ns |

Supporting Table 3: Effects of treatment, duration and intial food concentration on the mean number of fecal pellets produced during the experiments. The upper part of the table shows results from a three-way ANOVA of the effects of treatment, experimental duration (48h in Exp. 5B, 24h in the others) and intial food particle concentration (*high* in Exp. 5 and *low* in the others) on number of fecal pellets. 'SSn': Sum of Squares in the numerator (i.e. SS effect); 'SSd': Sum of Squares in the denominator (i.e. SS error); 'DFn': Degrees of Freedom effect; 'DFd': Degrees of Freedom error; 'F': F-value; 'p': p-value; 'p<.05': significance at the 0.05 level; 'ges': Generalized Eta-Squared measure of effect size.

The lower part of the table shows results from Tukey HSD *post hoc* assessing the significance of differences between pairs of treatments (the other two variables had only two levels). 'n1'/'n2': the number of observations in group1 and group2; 'estimate': difference in observed means in group1 and group2; 'conf.low'/'conf.high': lower and upper end points of the confidence interval around the difference; 'p.adj': p-value after adjustment for multiple comparisons; 'p.adj.signif': significance at the 0.05 level (ns: not significant). Food treatments: LAMHY-FE, LAMHY-FR: fresh or frozen *L. hyperborea*, respectively; SACLA-FE, SACLA-FR: fresh or frozen *S. latissima*, respectively; NONE: no food, PHYTO: phytoplankton (*R. baltica*).

| Effect | SSn | SSd | DFn | DFd | F | p | p<.05 | ges |
|---|---|---|---|---|---|---|---|---|
| TREATMENT | 5,122 | 246 | 5 | 18 | 75.1 | 2.0e-11 | * | 0.95 |
| DURATION | 151 | 246 | 1 | 18 | 11.1 | 4.0e-03 | * | 0.38 |
| INITIAL_CONCENTRATION | 34 | 246 | 1 | 18 | 2.5 | 1.3e-01 | | 0.12 |

| group1 | group2 | n1 | n2 | estimate | conf.low | conf.high | p.adj | p.adj.signif |
|---|---|---|---|---|---|---|---|---|
| LAMHY-FE | LAMHY-FR | 4 | 4 | 0.250 | -11.6 | 12.1 | 1.0e+00 | ns |
| LAMHY-FE | SACLA-FE | 4 | 7 | 5.071 | -5.4 | 15.6 | 6.6e-01 | ns |
| LAMHY-FE | SACLA-FR | 4 | 3 | 0.167 | -12.6 | 13.0 | 1.0e+00 | ns |
| LAMHY-FR | SACLA-FE | 4 | 7 | 4.821 | -5.7 | 15.3 | 7.0e-01 | ns |
| LAMHY-FR | SACLA-FR | 4 | 3 | -0.083 | -12.9 | 12.7 | 1.0e+00 | ns |
| NONE | PHYTO | 3 | 5 | 34.933 | 22.7 | 47.2 | 2.6e-07 | **** |
| NONE | LAMHY-FE | 3 | 4 | -2.167 | -15.0 | 10.6 | 9.9e-01 | ns |
| NONE | LAMHY-FR | 3 | 4 | -1.917 | -14.7 | 10.9 | 1.0e+00 | ns |
| NONE | SACLA-FE | 3 | 7 | 2.905 | -8.7 | 14.5 | 9.7e-01 | ns |
| NONE | SACLA-FR | 3 | 3 | -2.000 | -15.7 | 11.7 | 1.0e+00 | ns |
| PHYTO | LAMHY-FE | 5 | 4 | -37.100 | -48.3 | -25.9 | 2.3e-08 | **** |
| PHYTO | LAMHY-FR | 5 | 4 | -36.850 | -48.1 | -25.6 | 2.6e-08 | **** |
| PHYTO | SACLA-FE | 5 | 7 | -32.029 | -41.8 | -22.2 | 2.8e-08 | **** |
| PHYTO | SACLA-FR | 5 | 3 | -36.933 | -49.2 | -24.7 | 1.0e-07 | **** |
| SACLA-FE | SACLA-FR | 7 | 3 | -4.905 | -16.5 | 6.7 | 7.6e-01 | ns |

Supporting Table 4: Results from Tukey HSD *post hoc* tests comparing the mean estimated *S. latissima* DNA content produced for different treatments. n1'/'n2': the number of observations in group1 and group2; 'estimate': difference in observed means in group1 and group2; 'conf.low'/'conf.high': lower and upper end points of the confidence interval around the difference; 'p.adj': p-value after adjustment for multiple comparisons; 'p.adj.signif': significance at the 0.05 level (ns: not significant). Treatments: SACLA-FE, SACLA-FR: fresh or frozen *S. latissima*, respectively; NONE: no food, PHYTO: phytoplankton (*R. baltica*); SACLA-FE-SEDATED: copepod submerged in fresh *S. latissima* treatment while sedated; BLANK: No sample.

| group1 | group2 | n1 | n2 | estimate | conf.low | conf.high | p.adj | p.adj.signif |
|---|---|---|---|---|---|---|---|---|
| BLANK | NONE | 3 | 8 | 1.4e-17 | -0.064 | 0.064 | 1.0e+00 | ns |
| BLANK | PHYTO | 3 | 6 | -8.1e-18 | -0.066 | 0.066 | 1.0e+00 | ns |
| BLANK | SACLA-FE | 3 | 8 | 3.7e-02 | -0.027 | 0.101 | 4.8e-01 | ns |
| BLANK | SACLA-FR | 3 | 4 | 1.6e-01 | 0.089 | 0.232 | 5.2e-06 | **** |
| NONE | PHYTO | 8 | 6 | -2.2e-17 | -0.051 | 0.051 | 1.0e+00 | ns |
| NONE | SACLA-FE | 8 | 8 | 3.7e-02 | -0.010 | 0.084 | 1.8e-01 | ns |
| NONE | SACLA-FR | 8 | 4 | 1.6e-01 | 0.103 | 0.218 | 1.1e-07 | **** |
| PHYTO | SACLA-FE | 6 | 8 | 3.7e-02 | -0.014 | 0.088 | 2.5e-01 | ns |
| PHYTO | SACLA-FR | 6 | 4 | 1.6e-01 | 0.100 | 0.221 | 3.0e-07 | **** |
| SACLA-FE | SACLA-FR | 8 | 4 | 1.2e-01 | 0.066 | 0.181 | 1.0e-05 | **** |
| BLANK | SACLA-FE-SEDATED | 3 | 1 | 4.4e-02 | -0.065 | 0.152 | 8.1e-01 | ns |
| NONE | SACLA-FE-SEDATED | 8 | 1 | 4.4e-02 | -0.056 | 0.143 | 7.5e-01 | ns |
| PHYTO | SACLA-FE-SEDATED | 6 | 1 | 4.4e-02 | -0.058 | 0.145 | 7.6e-01 | ns |
| SACLA-FE | SACLA-FE-SEDATED | 8 | 1 | 6.7e-03 | -0.093 | 0.106 | 1.0e+00 | ns |
| SACLA-FR | SACLA-FE-SEDATED | 6 | 1 | -1.2e-01 | -0.222 | -0.012 | 2.3e-02 | * |

Supporting Table 5: Results from Tukey HSD *post hoc* tests comparing the mean estimated *L. hyperborea* DNA content produced for different treatments. 'n1'/'n2': the number of observations in group1 and group2; 'estimate': difference in observed means in group1 and group2; 'conf.low'/'conf.high': lower and upper end points of the confidence interval around the difference; 'p.adj': p-value after adjustment for multiple comparisons; 'p.adj.signif': significance at the 0.05 level (ns: not significant). Treatments: LAMHY-FE, LAMHY-FR: fresh or frozen *L. hyperborea*, respectively; NONE: no food, PHYTO: phytoplankton (*R. baltica*); LAMHY-FE-SEDATED: copepod submerged in fresh *L. hyperborea* treatment while sedated; BLANK: No sample.

| group1 | group2 | n1 | n2 | estimate | conf.low | conf.high | p.adj | p.adj.signif |
|---|---|---|---|---|---|---|---|---|
| BLANK | NONE | 3 | 8 | 1.1e-03 | -3.5e-02 | 0.0373 | 1.0000 | ns |
| BLANK | PHYTO | 3 | 6 | -7.2e-18 | -3.8e-02 | 0.0378 | 1.0000 | ns |
| BLANK | LAMHY-FE | 3 | 8 | 3.6e-02 | -8.7e-05 | 0.0722 | 0.0508 | ns |
| BLANK | LAMHY-FR | 3 | 8 | 4.3e-03 | -3.2e-02 | 0.0404 | 0.9990 | ns |
| LAMHY-FE | LAMHY-FR | 8 | 8 | -3.2e-02 | -5.8e-02 | -0.0051 | 0.0127 | * |
| NONE | PHYTO | 8 | 6 | -1.1e-03 | -3.0e-02 | 0.0277 | 1.0000 | ns |
| NONE | LAMHY-FE | 8 | 8 | 3.5e-02 | 8.2e-03 | 0.0616 | 0.0051 | ** |
| NONE | LAMHY-FR | 8 | 8 | 3.1e-03 | -2.4e-02 | 0.0299 | 0.9990 | ns |
| PHYTO | LAMHY-FE | 6 | 8 | 3.6e-02 | 7.2e-03 | 0.0649 | 0.0080 | ** |
| PHYTO | LAMHY-FR | 6 | 8 | 4.3e-03 | -2.5e-02 | 0.0331 | 0.9970 | ns |
| BLANK | LAMHY-FE-SEDATED | 3 | 1 | 7.3e-02 | 1.2e-02 | 0.1349 | 0.0130 | * |
| LAMHY-FE | LAMHY-FE-SEDATED | 8 | 1 | 3.7e-02 | -2.0e-02 | 0.0938 | 0.3660 | ns |
| LAMHY-FR | LAMHY-FE-SEDATED | 8 | 1 | 6.9e-02 | 1.2e-02 | 0.1256 | 0.0104 | * |
| NONE | LAMHY-FE-SEDATED | 8 | 1 | 7.2e-02 | 1.5e-02 | 0.1287 | 0.0068 | ** |
| PHYTO | LAMHY-FE-SEDATED | 6 | 1 | 7.3e-02 | 1.6e-02 | 0.1309 | 0.0069 | ** |

Supporting Table 6: Species-specific primer pairs used in the quantitative polymerase chain reaction (qPCR) assays targeting *Laminaria hyperborea*, *Saccharina latissima* and *Calanus finmarchicus*. A 2-step amplification protocol was applied with an initial denaturation step at 98°C for 2 min followed by 45 cycles, 5s 98°C denaturation and 20s elongation at annealing temperature 62°C for L. hyperborea and S. latissima, and 54°C for C. finmarchicus. We used a total reaction volume of 15 µl, with 7.5 µl SsoFast EvaGreen Supermix (Bio-Rad, Hercules, CA, USA), 0.75 µl of each primer (0.5 µM final concentration), 4.5 µl sterilized pure water and 1.5 µL sample (individual copepods extracted in 0.2 ml QE). COI: Cytochrome oxidase subunit-1; Ta.°C: annealing temperature; bp: produced amplicon (base pair). References are: [1], [2], [3] and Anglès d'Auriac et al., in press. The annealing temperature for the *C. finmarchicus* primer was modified in this study.

| Target | Specificity | Primer name | Sequence | Ta.°C | bp | Ref |
|---|---|---|---|---|---|---|
| COI | *Laminaria hyperborea* | L_hyper-coi471F20 | CTCCCGGTATGACAATGGAT | 62 | 88 | [1,2] |
| | | L_hyper-coi538R21 | AAAACAGGAAGCGATAACAGT | | | |
| COI | *Saccharina latissima* | S_lat-coi286F22 | GCTTCTAGCGTCCTCATTGGTA | 62 | 106 | Anglès d'Auriac |
| | | S_lat-coi391R22 | AGCTAAGTCAACTGAAGGTCCT | | | |
| COI | *Calanus finmarchicus* | COI-2011 | YTCATCACTGCTGTCCTC | 54 | 117 | [3] |
| | | COI-2128R | GTGCTGRTATAAAATAGG | | | |

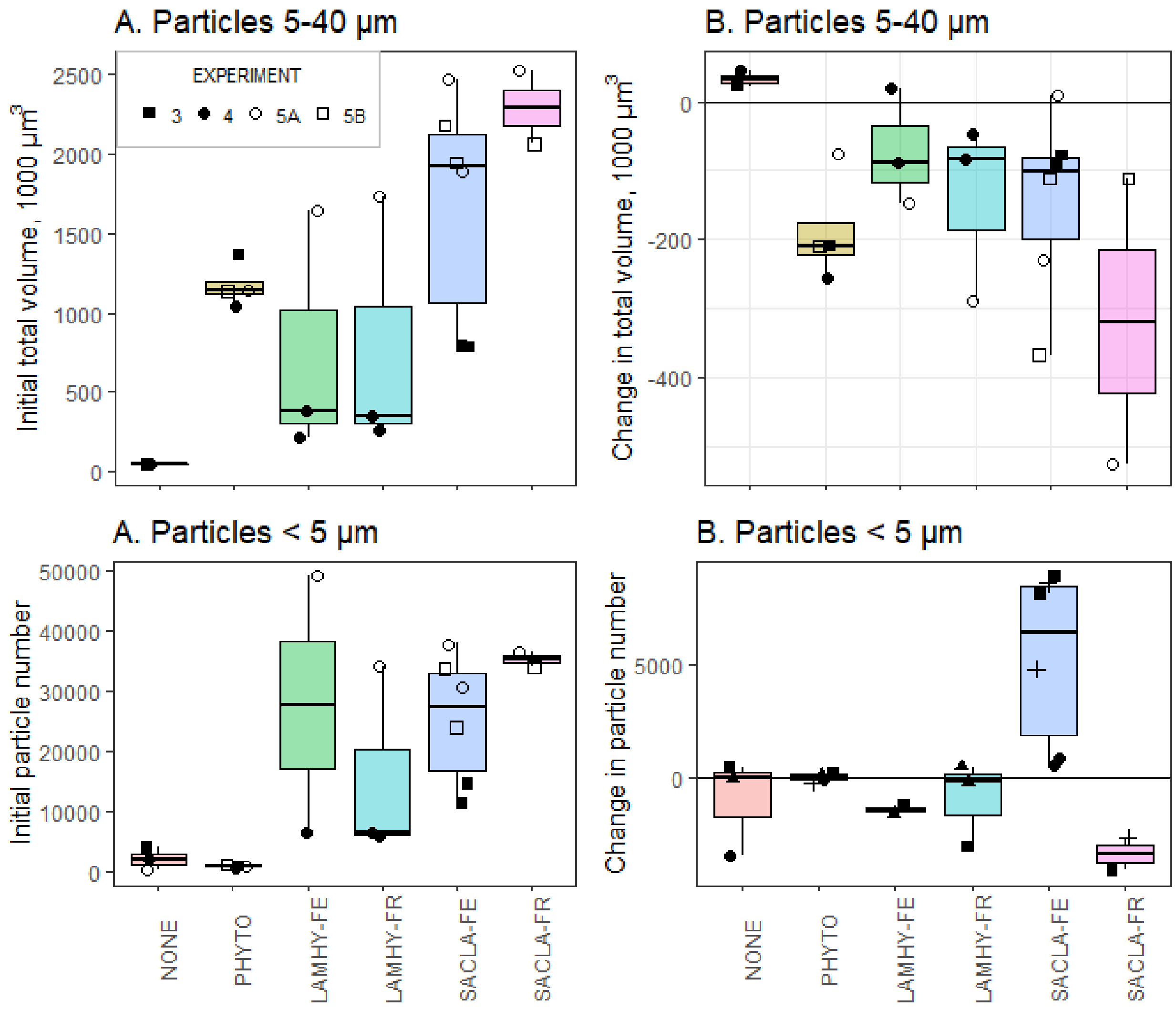


*Supporting Figure 1: Initial total volume of particles in the water (A) and change in total volume during the experiments (B) for particles 5-40 µm, the normal feeding range of* C. finmarchicus, *and initial number of particles in the water (C) and change in particle concentration during the experiments (D) for particles in the size range < 5µm, which is smaller than the normal feeding range of* C. finmarchicus. *Each symbol represents one observation (i.e., one experimental bottle measured before and after the experiment), with n per treatment ranging between 3 and 6, see Supplementary Table 3 for details. The overlaying box plots show the median (line), the interquartile range (box), and 1.5 × the interquartile range (whiskers) per treatment. Unfilled symbol are observations from Exp. 5 with a higher initial volume of kelp than the other experiments. Experiments 3-4 lasted 24 h, while Exp. 5 lasted 48 h. But also here, the water was changed and the number of particles counted after each 24 h interval (Exp. 5A and 5B, respectively). Particle count data are lacking for Exp. 1-2. Food treatments: NONE: no food, PHYTO: phytoplankton (*R. baltica*); LAMHY-FE, LAMHY-FR: fresh or frozen* L. hyperborea, *respectively; SACLA-FE, SACLA-FR: fresh or frozen* S. latissima, *respectively.*

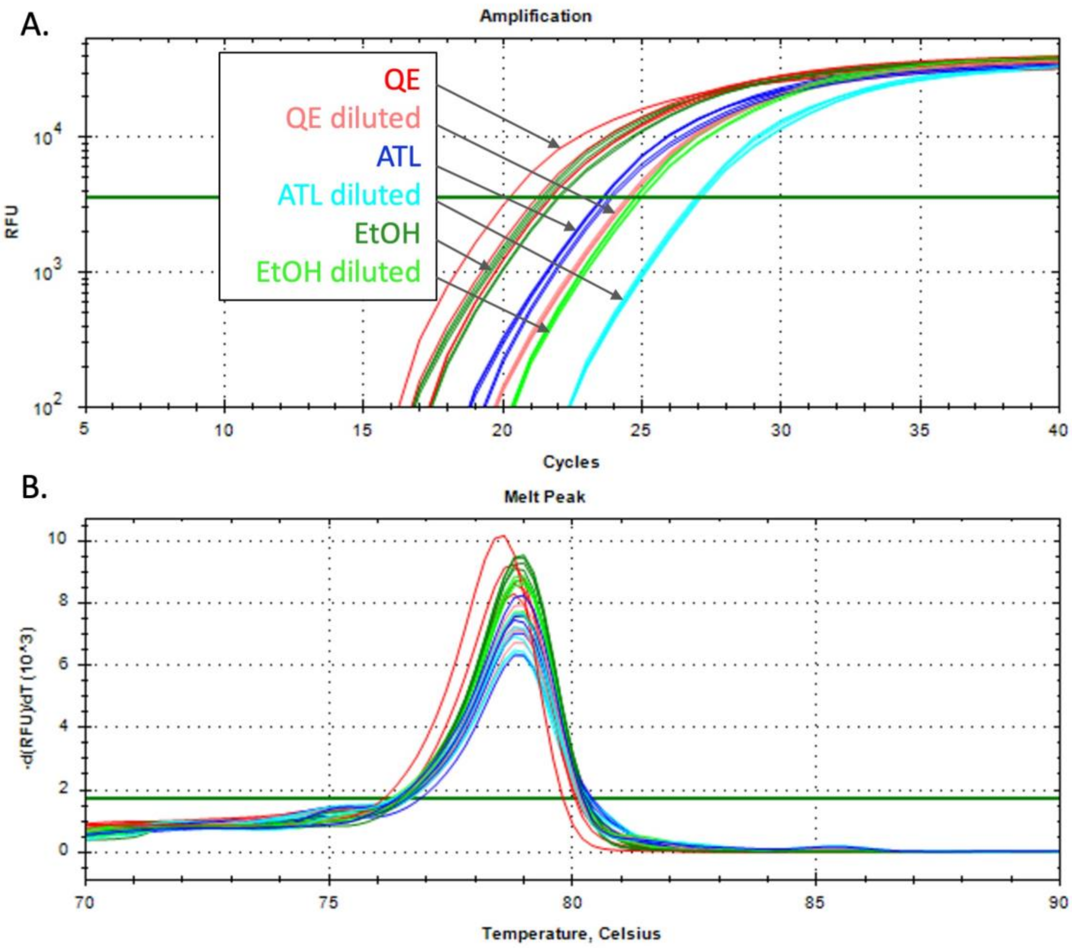


*Supporting Figure 2: Test results for the* C. finmarchicus *species-specific gradient (45.0-60.0°C) qPCR for from copepod samples preserved in different solutions (ethanol, ATL buffer or QE), showing strong and specific detection when applying an annealing temperature less than 54.5°C, with strongest detection using QE (red lines). DNA amplification was determined for samples extracted as described in the Method section and after a 10 times dilution in sterile water.*
